# Peculiar hybrid genomes of devastating plant pests promote plasticity in the absence of sex and meiosis

**DOI:** 10.1101/046805

**Authors:** Romain Blanc-Mathieu, Laetitia Perfus-Barbeoch, Jean-Marc Aury, Martine Da Rocha, Jérôme Gouzy, Erika Sallet, Cristina Martin-Jimenez, Philippe Castagnone-Sereno, Jean-François Flot, Djampa K Kozlowski, Julie Cazareth, Arnaud Couloux, Corinne Da Silva, Julie Guy, Corinne Rancurel, Thomas Schiex, Pierre Abad, Patrick Wincker, Etienne G.J. Danchin

**Affiliations:** INRA, Univ. Nice Sophia Antipolis, CNRS, UMR 1355-7254 Institut Sophia Agrobiotech, 06900 Sophia Antipolis, France.; Commissariat à l’Energie Atomique (CEA), Institut de Génomique (IG), Genoscope, Evry, BP5706, 91057, France.; INRA, CNRS, UMR 441-2594, Laboratoire des Interactions Plantes-Microorganismes (LIPM), Castanet-Tolosan, F-31326, France; Université Libre de Bruxelles (ULB), Evolutionary Biology & Ecology, C.P. 160/12, Avenue F.D. Roosevelt 50, 1050 Brussels, Belgium.; CNRS, Univ. Nice Sophia Antipolis, Institute of Molecular and Cellular Pharmacology, 06560 Sophia Antipolis, France; INRA, Unité de Mathématiques et Informatique UR 875, F-31320 Castanet Tolosan, France; Université d’Evry Val d’Essonne, UMR 8030, Evry, CP5706, 91057, France; Centre National de Recherche Scientifique (CNRS), UMR 8030, Evry, CP5706, 91057, France

## Abstract

Root-knot nematodes (genus *Meloidogyne*) show an intriguing diversity of reproductive modes ranging from obligatory sexual to fully asexual reproduction. Intriguingly, the most damaging species to the world agriculture are those that reproduce without meiosis and without sex. To understand this parasitic success despite the absence of sex and genetic exchanges, we have sequenced and assembled the genomes of 3 obligatory ameiotic asexual *Meloidogyne* species and have compared them to those of meiotic relatives with facultative or obligatory asexual reproduction. Our comparative genomic analysis shows that obligatory asexual root-knot nematodes have a higher abundance of transposable elements (TE) compared to the facultative sexual and contain duplicated regions with a high within-species average nucleotide divergence of 8%. Phylogenomic analysis of the genes present in these duplicated regions suggests that they originated from multiple hybridization events. The average nucleotide divergence in the coding portions between duplicated regions is ~5-6 % and we detected diversifying selection between the corresponding gene copies. Genes under diversifying selection covered a wide spectrum of predicted functional categories which suggests a high impact of the genome structure at the functional level. Contrasting with high within-species nuclear genome divergence, mitochondrial genome divergence between the three ameiotic asexuals was very low, suggesting that these putative hybrids share a recent common maternal donor lineage. The intriguing parasitic success of mitotic root-knot nematodes in the absence of sex may be partly explained by TE-rich composite genomes resulting from multiple allo-polyploidization events and promoting plasticity in the absence of sex.

## INTRODUCTION

Fully asexual reproduction (i.e. obligate parthenogenesis) is estimated to occur in only ~0.1% of animal lineages, which generally occupy shallow branches in the animal tree of life (Vrijenhoek 1998; Rice 2002). Although there are some exceptions (Neiman et al. 2009; Danchin et al. 2011; Flot et al. 2013), the majority of asexual lineages of animals seem to be recently derived from sexual lineages, suggesting that asexual lineages are generally short-lived. Asexual animals lack the possibility to combine advantageous alleles from different individuals via sexual recombination. Furthermore, Muller’s ratchet (Muller 1964) and Kondrashov’s hatchet (Kondrashov 1988) models of “clonal decay” predict that they progressively accumulate deleterious mutations. Supporting these models, different studies have demonstrated accelerated accumulation of harmful mutations in asexual lineages (Paland and Lynch 2006; Barraclough et al. 2007; Neiman et al. 2010; Henry et al. 2012; Hollister et al. 2015), or increased accumulation of transposable elements in the absence of sex (Schaack et al. 2010; Ågren et al. 2015). Hence, it is commonly postulated that obligate parthenogenetic animals have evolutionary and adaptive disadvantages compared to their sexual relatives and therefore represent evolutionary dead ends. According to the geographical parthenogenesis model, parthenogenetic populations of plant and animals are generally present at the edge of the geographical distribution of species, in marginal or anthropologically disturbed environments (Hörandl 2009; Vrijenhoek and Parker 2009). Their uniparental clonal reproductive mode is supposed to be advantageous for colonizing marginal environments where they escape competition with their sexual relatives. Although this last statement is controversial in the literature, parthenogenetic species are indeed frequently found at higher latitudes and altitudes (Vrijenhoek and Parker 2009).

Root-knot nematodes (genus *Meloidogyne*) display a variety of reproductive modes ranging from classical sexual reproduction (amphimixis) to obligate asexual reproduction (apomixis) with intermediates able to alternate between sexual (amphimixis) and asexual (automixis) reproduction (Castagnone-Sereno et al. 2013). These notorious plant pests have been ranked number one in terms of economic threat to the agriculture among all plant-parasitic nematodes (Jones et al. 2013). Challenging the view that fully asexual lineages of animals are outcompeted by their sexual relatives, *Meloidogyne* species that reproduce without meiosis and without sex have a broader host range, a wider and more southern geographical distribution and are more devastating than their sexual relatives (Castagnone-Sereno 2006; Castagnone-Sereno and Danchin 2014). Whether some genomic singularities could underlie the higher parasitic success of asexually reproducing *Meloidogyne* remains unclear. In 2008, we coordinated the publication of the draft genome sequence of *Meloidogyne incognita* (Abad et al. 2008), an obligatory asexual nematode and the draft genome of *Meloidogyne hapla*, a facultative sexual, was published the same year (Opperman et al. 2008). One singularity of the *M. incognita* genome was the presence of genomic regions in two or more copies that spanned several megabases of the genome and had an average nucleotide divergence of ~8% (Abad et al. 2008). Such a structure was not identified in the facultative sexual *M. hapla* and a few examples of duplicated genomic regions in *M. incognita* were searched in *M. hapla* but identified in one single copy (Bird et al. 2009). More recently, a draft genome of the obligatory meiotic asexual *M. floridensis* has been published (Lunt et al. 2014). Although the high degree of fragmentation of this assembly prevented identification of duplicated genomic regions, a huge proportion of protein-coding genes was found to be duplicated and suggested a possible duplicated structure too.

The possible origin of the duplicated and divergent genomic regions observed in apomictic *Meloidogyne* as well as their potential functional consequences remained so far unexplored at whole-genome level. The two main hypotheses for the origin of this duplicated genome structure are (i) that duplicated regions represent former paternal and maternal genomes that diverged and became rearranged after their diploid sexual ancestor became asexual or (ii) that they result from interspecific hybridization events (Castagnone-Sereno and Danchin 2014). The potential functional plasticity conferred by the duplicated genome structure of *M. incognita* has so far never been assessed. Furthermore, in the absence genomes for other apomictic *Meloidogyne*, it was impossible to evaluate whether such a duplicated genome structure is a specificity of *M. incognita* or a more general signature of root-knot nematodes with mitotic asexual reproduction.

Here, we have re-sequenced the genome of *M. incognita* at a much deeper coverage and have sequenced *de novo* the genomes of *M. javanica* and *M. arenaria*, two other apomictic root-knot nematodes. We have assembled these 3 genomes and validated genome assembly sizes with experimental assays. We have annotated the protein-coding genes and transposable elements (TE) of the 3 genomes and performed a comparative genomics analysis, including the genome of the facultative sexual species *M. hapla* and the meiotic parthenogenetic *M. floridensis*. We have compared the relative abundance of TE to determine whether they have undergone expansions / reductions in asexual compared to sexual *Meloidogyne*. Using a phylogenomic analysis on duplicated genomic regions conserved between species, we deciphered the origin and evolutionary history of the peculiar genome architecture of mitotic parthenogenetic *Meloidogyne*. To assess the potential functional outcome of the presence of duplicated genomic regions, we systematically searched signs of positive and diversifying selection in gene copies present in these genomic blocks at a whole-genome scale.

## RESULTS

### Genome assemblies are experimentally supported and show high completeness

We assembled the genomes of clonal lineages of 3 asexually reproducing *Meloidogyne* species. Our assembly pipeline (Methods) produced genome sequences with sizes of 184, 236 and 258 Mb, for *M. incognita* (*Mi*), *M. javanica* (*Mj*) and *M. arenaria* (*Ma*) respectively (Table 1). We measured DNA content via flow cytometry experiments and obtained genome size estimates of 189 ±15, 297 ±27 and 304 ±9 for *Mi*, *Mj* and *Ma*, respectively. The genome assembly size of *Mi* was in the range of estimated size via flow cytometry whereas *Mj* and *Ma* assemblies were smaller by 34-60 Mb. To check whether these differences in sizes could be explained by duplicated or repetitive regions collapsed during genome assembly, we plotted the distribution of read coverage along the genome (Methods, Supplementary Fig. S1). We found that 17.1 (Mi), 42.9 (Mj) and 21.6 (Ma) Mb of genomes assemblies have a coverage twice higher than the rest of the genome sequence and might represent collapsed repeated regions. Hence part of the differences sizes can be explained by these collapsed regions, as previously observed in the genome of the obligate mitotic rotifer *Adineta vaga* (Flot et al. 2013). The genome assemblies contained 97% (*Mi*), 96% (*Mj*) and 95% (*Ma*) of the 248 Core Eukaryotic Genes (CEG) in complete length (Parra et al. 2009). These are the highest scores for a *Meloidogyne* genome so far and suggest that the 3 mitotic parthenogenetic *Meloidogyne* genomes we have assembled are the most complete available to date. We annotated 45,351 (*Mi*), 98,578 (*Mj*) and 103,269 (*Ma*) genes (including, protein-coding genes, ncRNAs, rRNAs and tRNAs). Protein-coding sequences spanned up to 43.7 (24% *Mi*), 75.2 (32% *Mj*) and 82.2 (32% *Ma*) Mb of the genome sequences. Transposable elements (TE) covered 50.0 % (Mi), 50.8 % (Mj) and 50.8 % (Ma) of the genome assemblies. In comparison, only 29.2% of the *M. hapla* genome was covered by TE, using the exact same annotation protocol (Table 2). Due to its high fragmentation state, the genome of *M. floridensis* could not be annotated for TE. On average, 60% of the genes of mitotic parthenogenetic species overlap with or are embedded within TE, whereas this proportion reaches only 40% in *M. hapla* (see TE section for more details).

**Table 1.** Size, assembly and gene annotation statistics of *Meloidogyne* genomes

| Species | *M. incognita* * | *M. javanica* * | *M. arenaria* * | *M. hapla* | *M. floridensis* |
| --- | --- | --- | --- | --- | --- |
| Flow cytometry* (Mb) | 189 ±15 | 297 ±27 | 304 ±9 | NA | NA |
| Assembly size (Mb) | 183.53 | 235.80 | 258.07 | 53.60 | 99.89 |
| N50 (bp) | 38,588 | 10,388 | 16,462 | 83,645 | 3,516 |
| Complete CEG | 97% | 96% | 95% | 93% | 60% |
| # genes** | 45,351 | 98,578 | 103,001 | NA | NA |
| # CDS | 43,718 | 97,208 | 101,269 | 14,207 | 49,941 |
| Protein-coding (Mb) (% assembly) | 43.7 (23,8 %) | 75.2 (31,9 %) | 82.2 (32,1 %) | NA | NA |

\* This study. \*\* Including non-protein-coding genes NA= information not available

**Table 2.** Abundance and diversity of transposable elements in *Meloidogyne* genomes

| % genome / Species | *M. incognita* | *M. javanica* | *M. arenaria* | *M. hapla* |
| --- | --- | --- | --- | --- |
| **Class I:** | **18.05** | **18.92** | **19.20** | **12.11** |
| LINE: (copy#) | 1.57 (2067) | 1.52 (3012) | 1.53 (3214) | 1.31 (513) |
| LTR: (copy#) | 2.93 (3458) | 3.49 (5831) | 3.6 (6213) | 1.84 (677) |
| PLE: (copy#) | 0.03 (53) | 0.04 (86) | 0.03 (80) | 0.05 (13) |
| SINE: (copy#) | <0.01 (30) | <0.01 (50) | <0.01 (37) | <0.01 (5) |
| DIRS: (copy#) | 0.21 (221) | 0.29 (412) | 0.32 (460) | 0.05 (17) |
| Uncl.: (copy#) | 13.3 (25347) | 13.57 (37451) | 13.71 (39449) | 8.86 (5025) |
| **Class II:** | **10.53** | **10.02** | **10.34** | **5.39** |
| Helitron : (copy#) | 2.05 (1588) | 2.13 (2774) | 2.11 (2755) | 1.77 (525) |
| Maverick: (copy#) | 3.62 (100) | 3.31 (121) | 3.51 (139) | 1.54 (2) |
| TIR: (copy#) | 3.69 (7128) | 3.53 (9675) | 3.58 (10547) | 1.66 (977) |
| Uncl.: (copy#) | 1.17 (3577) | 1.04 (4478) | 1.13 (5169) | 0.43 (363) |
| **Other: cov.** | **21.41** | **21.82** | **21.24** | **11.65** |
| **Total:** | **49.99** | **50.76** | **50.78** | **29.16** |

### The genomes of asexual *Meloidogyne* are highly duplicated and gene copies form divergent syntenic blocks

We investigated the structure of *Meloidogyne* genomes by describing the duplication relationships of their protein-coding genes using MCScanX (Wang et al. 2012). MCScanX classifies protein-coding genes as (*i*) singleton when no duplicates are found in the assembly, (*ii*) proximal when duplicates are on the same scaffold and separated by 1 to 10 genes, (*iii*) tandem when duplicates are next to each other on the same scaffold, (*iv*) whole genome duplication (WGD) or segmental when duplicates form collinear blocks with other pairs of duplicated genes and (*v*) dispersed when the duplicates cannot be assigned to any of the other categories. According to MCScanX, the vast majority of the protein-coding genes in the obligatory asexual *Meloidogyne* species were duplicated: *Mi*: 40,675 (93.0%), *Mj*: 91,336 (94.0%) and *Ma*: 95,332 (94.1%)) vs. 6,625 (46.6%) for the facultative sexual *M. hapla* and 26,428 (52.9%) for the meiotic parthenogenetic *M. floridensis* (Table 3). MCScanX synteny analysis classified 12,445, 5,806 and 15,632 genes in the WGD / segmental category for the genomes of the ameiotic species *M. incognita*, *M. javanica* and *M. arenaria*, respectively. These protein-coding genes allowed us to define 933 (*Mi*), 581 (*Mj*) and 1,648 (*Ma*) pairs of duplicated genome regions. In contrast, there were only 90 segmentally duplicated genes (forming 11 pairs of duplicated regions) in the facultative sexual meiotic species *M. hapla* and even less (12 genes, 1 pair of regions) in the meiotic parthenogenetic *M. floridensis* (Table 4). Thus, the high incidence of duplicated genes forming collinear genomic regions appear specific to the ameiotic *Meloidogyne* species, although the low contiguity of the *M. floridensis* genome (N50=3.5kb) probably prevents identification of segmental duplications. Genomic sequence sizes of collinear regions sum up to 58.6, 14.8 and 59.0 Mb for *M. incognita*, *M. javanica* and *M. arenaria*, respectively corresponding to 31.8%, 6.3% and 23.0 % of the size of their respective genome assemblies (Table 4). Average nucleotide divergence between pairs of duplicated genomic regions was 8.4%, 7.5% and 8.2% for *Mi*, *Mj* and *Ma*, respectively, indicating a similar average divergence of ~8% between region pairs. The divergence levels were substantially lower in coding regions (4.7, 6.0 and 5.9%) than in non-coding regions (9.7, 9.0 and 9.7 % for intergenic and 11, 10.4 and 11.1% for introns) for *Mi*, *Mj* and *Ma*, respectively (Table 4, Supplementary Fig. S2).

**Table 3.** MCScanX classification of protein-coding genes in *Meloidogyne* genome sequences

| Species | *M. incognita* | *M. javanica* | *M. arenaria* | *M. hapla* | *M. floridensis* |
| --- | --- | --- | --- | --- | --- |
| Total CDS | 43,718 | 97,208 | 101,269 | 14,207 | 49,941 |
| Singleton | 3,043 (7.0%) | 5,872 (6.0%) | 5,937 (5.9%) | 7,582 (53.4%) | 23,513 (47.1%) |
| Dispersed | 25,964 (59.4%) | 78,989 (81.2%) | 73,569 (72.7%) | 4,891 (34.4%) | 25,610 (51.3%) |
| Proximal | 1,026 (2.4%) | 2,116 (2.2%) | 2,647 (2.6%) | 791 (5.6%) | 168 (0.3%) |
| Tandem | 1,240 (2.8%) | 4,425 (4.6%) | 3,484 (3.4%) | 853 (6.0%) | 638 (1.3%) |
| WGD / segmental | 12,445 (28.5%) | 5,806 (6.0 %) | 15,632 (15.4%) | 90 (0.6%) | 12 (~0%) |
| N50 (kb) | 38.6 | 10.4 | 16.5 | 83.6 | 3.5 |

**Table 4.** Number of pairs of duplicated regions, cumulative size and divergence

| Species |  | *M. incognita* | *M. javanica* | *M. arenaria* |
| --- | --- | --- | --- | --- |
| # pairs of duplicated regions |  | 933 | 581 | 1648 |
| Cumulative size of duplicated collinear regions(Mb) (% genome sequence) |  | 58.6 (31.8%) | 14.8 (6.3%) | 59.0 (23.0%) |
| % nucleotide divergence (standard deviation) computed from NUCmer alignments | Overall | 8.4 (2.5) | 7.5 (2.9) | 8.2 (2.7) |
|  | intergenic | 9.7 (3.3) | 9.0 (4.2) | 9.7 (3.5) |
|  | introns | 11.0 (2.8) | 10.4 (3.2) | 11.1 (2.4) |
|  | CDS | 4.7 (1.7) | 6.0 (2.7) | 5.9 (2.4) |
| Median CDS % divergence (MCscanX) |  | 3.5 | 3.8 | 4.1 |
| Median protein % divergence (MCscanX) |  | 2.7 | 3.7 | 4 |
| Median Ks |  | 0.1 | 0.1 | 0.1 |

Median rates of synonymous substitutions (Ks) between gene pairs forming duplicated regions were 0.1 for the three apomictic Meloidogyne (as measured using the Nei and Gojobori method (Nei and Gojobori 1986) implemented in MCScanX). Ks averaged for duplicated pairs of regions were significantly (P<10^−8^ for *Mi*, *Mj* and *Ma*) negatively correlated with collinearity, which we measured as the fraction of collinear genes within a region pair (Supplementary Fig. S3). This indicates that divergence in terms of number of conserved genes between a pair of genome regions correlates with the nucleotide divergence of their coding sequences. In *M. incognita* and *M. arenaria*, we found 5 instances of duplicated regions on a same scaffold (Fig. 1). In *M. incognita*, this corresponded to 42 pairs of anchor genes present in 4 pairs of tandem regions and 1 palindrome, whereas in *M. arenaria*, we found 29 pairs of anchor genes present in 2 pairs of tandem regions and 3 palindromes. As in the genome of *A. vaga* (Flot et al. 2013), such structures appear incompatible with chromosome pairing during meiosis if we consider that the duplicated regions represent vestiges of homologous chromosomes. No similar structure was found in the *M. javanica* nor in the *M. hapla* or *M. floridensis* genomes. Average Ks value of gene pairs forming tandem or palindromic regions were in the range of Ks values measured for gene pairs in the rest of duplicated regions, suggesting they have the same divergence times.

**Fig 1.**
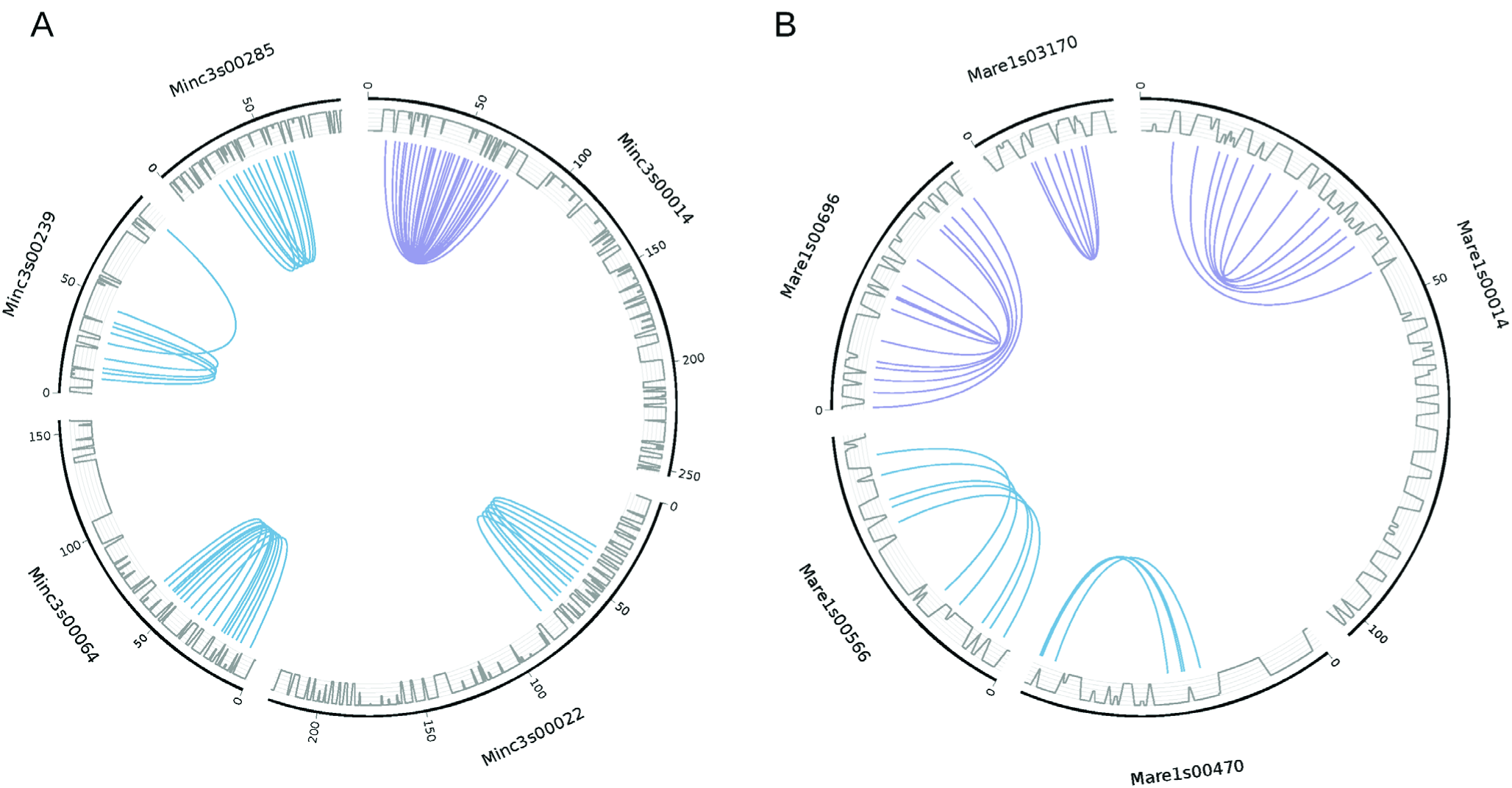
A genome structure incompatible with conventional meiosis. Five pairs of duplicated collinear regions are found to occur on a same scaffold for *M. incognita* (A) and five for *M. arenaria* (B). All curves show connections between anchor gene pairs used by MCScanX to define duplicated regions (blue curves show tandem duplications and purple curves show palindromic duplications). Grey lines represent the gene density on the scaffolds.

In plant genomes, following WGD, a fragmentation bias has been observed. One genomic copy tends to retain more genes and accumulate less mutations than the other copy (Murat et al. 2014). For each pair of collinear regions in *Meloidogyne*, we tested whether a bias of retention of ancestral genes could be observed between the two regions (Methods). We found 23 (*Mi*), 2 (*Mj*) and 37 (*Ma*) cases where one region had significantly (Chi-square test, 5% level) lost more ancestral genes than its counterpart.

### The duplicated regions have different origins and evolutionary histories

To decipher the origin and evolutionary history of the duplicated structure observed in the mitotic parthenogenetic *Meloidogyne*, we conducted a phylogenomic analysis focused on the gene copies forming pairs of duplicated regions. We searched for the most frequent topologies among phylogenies performed on the anchor genes pairs used to define synteny blocks conserved between the three apomictic *Meloidogyne* species and with the amphimictic *M. hapla* set as a reference outgroup. We used a dataset composed of 60 groups of homologous genomic regions defined as follows. The genomic regions must contain at least 3 collinear genes in 2 copies in each of the apomictic *Meloidogyne* vs. one single copy in the amphimictic *M. hapla* (Fig. 2.A for an example). These 60 groups of genomic regions encompass 2,202 orthologous and paralogous genes distributed in 222 clusters of orthologs (Fig. 2.B for an example, Supplementary Fig. S4 for the 60 shared homologous regions). Within each of the 60 groups of genomic regions, multiple alignments of all the clusters of homologous genes were generated individually. Although duplicated genes within a species are, on average, relatively distant (Ks=0.1, 5-6% nucleotide divergence) gene copies between species can occasionally be nearly identical. In order to maximize phylogenetic signal, the multiple alignments obtained for all the genes for each group of conserved regions were concatenated. From the 60 concatenated multiple alignments, we successfully generated 54 maximum-likelihood (ML) phylogenies (6 failed because of short alignments, see Fig. 2C for an example and Supplementary Fig. S4 for the 54 ML trees of conserved homologous regions).

**Fig 2.**
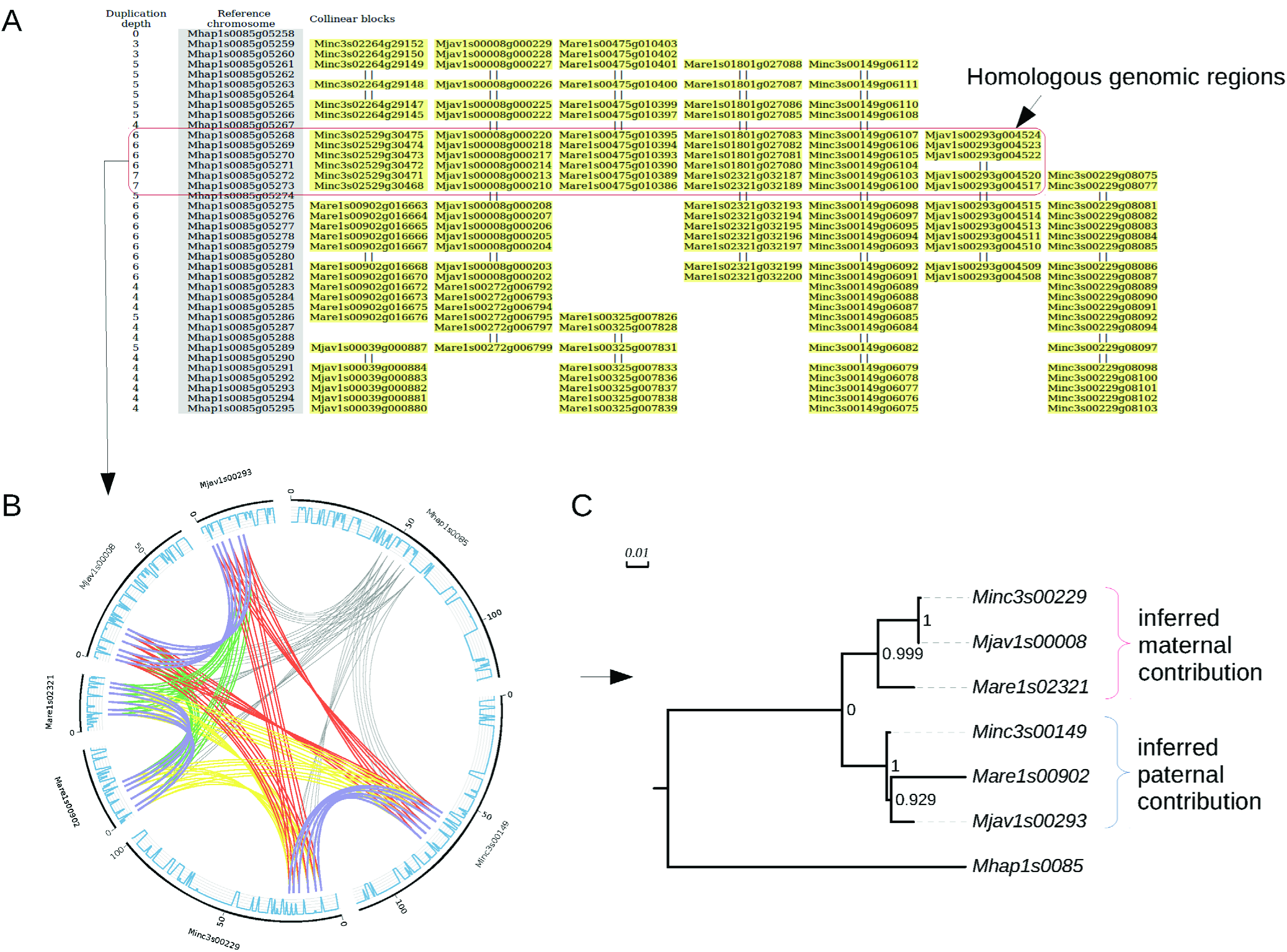
Structural and evolutionary relationship between pairs of duplicated regions for one of the 60 regions shared by all species. **A.** MCScanX output synthesizing the collinearity within and between *Meloidogyne* species. The facultative sexual species *M. hapla* was set as the reference. These files were parsed to select anchor genes pairs used in phylogenetic analyses (see method). **B.** Circos (Krzywinski et al. 2009) plot showing the anchor genes pairs (forming homologous regions) that were used for phylogenetic analyses. All curves show the connections between the anchor gene pairs used by MCScanX to detect blocks. In each circus plot, color codes are as follows. Collinear orthologs between *M. hapla* and any of the three asexuals species are in grey. Collinear ‘homeologs’ within asexual species are in purple. Collinear orthologs between *M. arenaria* and *M. javanica* are in green. Collinear orthologs between *M. arenaria* and *M. incognita* are in yellow. Collinear orthologs between *M. incognita* and *M. javanica* are in red. The outer blue lines represent the gene density on the scaffolds. **C.** Maximum-likelihood phylogeny of concatenated alignments of anchor protein-coding genes used to form blocks with SH-like branch support.

Among the 54 ML trees we generated, the vast majority (40 trees) presented at least 1 monophyletic clade made of at least 1 copy from each apomictic *Meloidogyne* species (Fig. 3). Only three possible bifurcating tree topologies exist to separate these clades: (1): (*Mi*, (*Ma*; *Mj*)), (2): (*Ma*, (*Mi*; *Mj)*) or (3): (*Mj*, (*Ma*; *Mi*)). Among all the possible clade topologies the most frequent was topology 1, observed 33 times. Topology 2 was observed 15 times and topology 3, 12 times (Fig. 3). A total of 20 trees combined two of the three clade topologies mentioned above and allowed testing whether the two duplicated regions presented the same evolutionary history. Among these 20 trees, a majority (13) contained clades with two different topologies, suggesting that these two regions had different evolutionary histories (Fig.4). The combination of topologies 1 and 2 was observed 7 times (see Fig. 2C for an example). The combination of topologies 1 and 3 and the combination of topologies 2 and 3 were each observed 3 times. Only 7 of the 20 trees showed twice the same topology; and in all these cases this was twice topology 1. Part of the genes forming segmental duplications were present in more than two copies in at least one apomictic species. We identified 387 groups of homologous anchors genes (total of 4,262 genes) for which at least one apomictic *Meloidogyne* species had 3 or 4 copies and the other two species had at least 2 copies. To decipher the evolutionary history of these additional copies, we counted the number of times the third or fourth copies hold a recent in-paralog (or allele-like position), relative to another copy vs. the number of times these copies were in a new independent branching position (Supplementary Fig. S5). For genes presents in 3 copies in a given genome assembly, the number of allele-like relationship was significantly lower (binomial test, P<10^−6^) than the number of new phylogenetic position, for all three species (Table 5). Hence, genes present in 3 copies more frequently formed a new independent branch in the phylogenetic trees than species-specific recent paralogs or allele-like branches. For genes in four copies within a given genome assembly, the number of allelic like relationship was significantly lower (binomial test, P<10^−5^) than the number of new positions, for *M. arenaria* only (Table 5).

**Fig 3.**
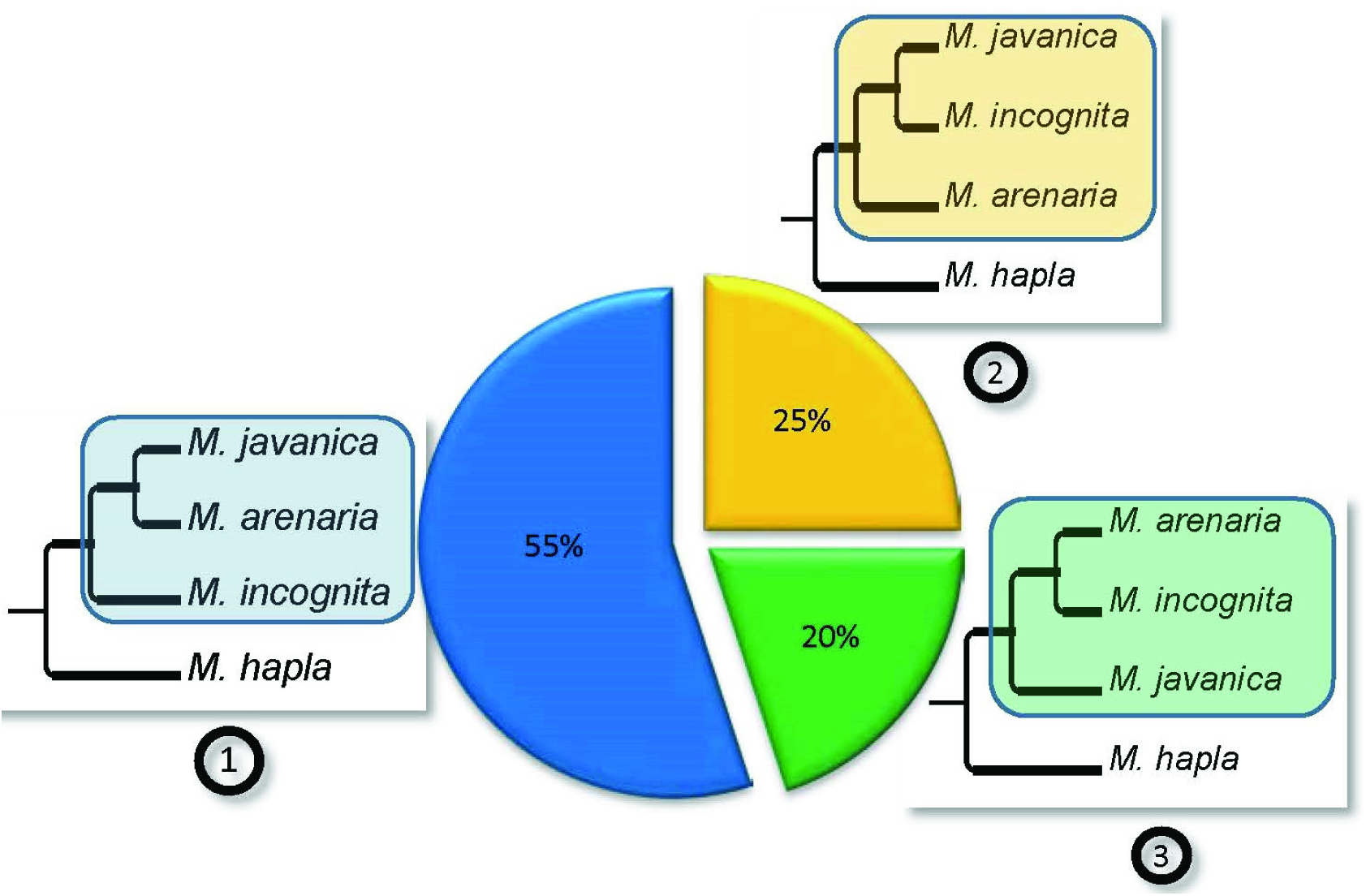
Phylogenetic relationships between duplicated regions in the genomes of apomictic *Meloidogyne*. The three possible topologies for bifurcating trees separating *M. incognita*, *M. javanica, M. arenaria* and their sexual relative *M. hapla* are represented as (1), (2) and (3) and their relative observed frequencies are indicated in the associated pie chart. The frequencies were calculated from the 40 phylogenetic trees containing at least one monophyletic clade with the 3 apomictic *Meloidogyne* that were constructed from the concatenated alignments covering a total of 2,202 protein-coding genes.

**Fig 4.**
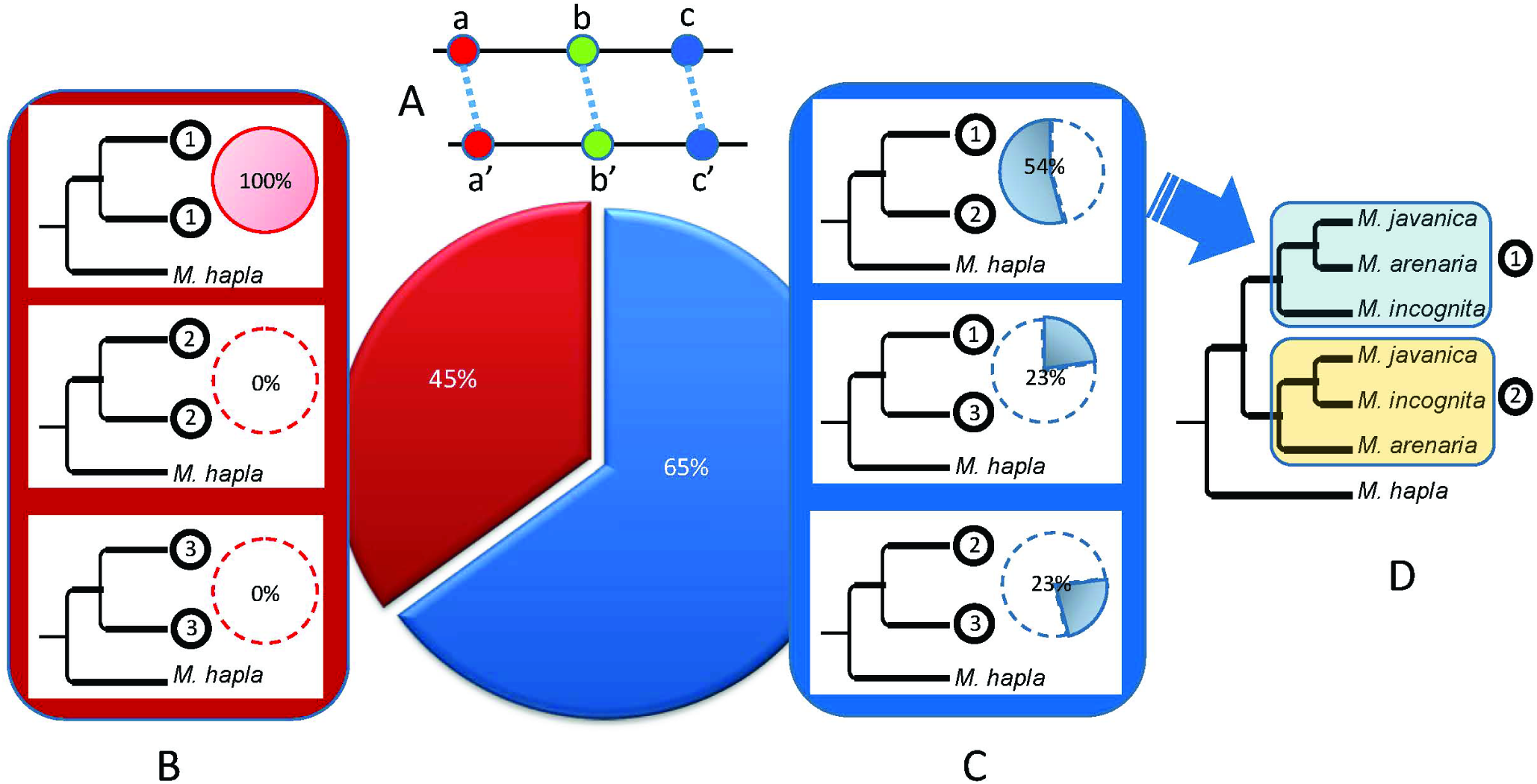
Combinations of topologies observed in trees combining at least two duplicated regions in the 3 parthenogenetic Meloidogyne. (A) Schematic representation of two duplicated regions each containing 3 collinear genes (a, b and c). (B) Topologies combining twice the same subclades among (1), (2) and (3), for the two duplicated regions (further detailed in Fig. 3). (C) Topologies combining two different subclades for the duplicated regions. The relative frequencies of trees combining twice the same (red) or two different (blue) subclades are indicated in the big central pie chart. Relative frequencies within the red and blue categories are indicated by small pie charts next to the corresponding schematic tree. (D) The most frequently observed topology is a combination of subclades (1) and (2).

**Table 5.** Number of trees supporting allelic-like *versus* homeologs evolutionary relationship for genes in regions present in 3 or 4 copies per species.

| Nb copies | Relation | *M. incognita* | *M. javanica* | *M. arenaria* |
| --- | --- | --- | --- | --- |
| 3 | Allele-like | 33 trees *** | 11 trees *** | 19 trees *** |
|  | Homeologs | 95 trees | 80 trees | 226 trees |
| 4 | Allele-like | 3 trees | 5 trees *NS* | 6 trees ** |
|  | Homeologs | 1 trees | 12 trees | 31 trees |

\*\*\*: $P < 10^{-6}$ , \*\*: $P < 10^{-3}$ , *NS* = non-significant, according to a binomial test

Overall, this ensemble of results suggests that the duplicated genome regions have different evolutionary histories and do not result from a common ancestral auto-polyploidisation.

### Asexual *Meloidogyne* share nearly identical mitochondrial genes

To reveal the maternal evolutionary history of *Meloidogyne* species included in our comparative genomics analysis, we performed a phylogeny based on mitochondrial protein-coding genes as well as the 12S and 16S mitochondrial rRNAs (Supplementary methods S1). The phylogenetic tree (Supplementary Fig. S6, Table S1) performed on the 12 concatenated protein-coding genes complemented by the rRNAs returned the following highly-supported topology: (*Ma*,((*Mi,Mf*),*Mj*))). This topology corresponds to clade topology 2, the second most frequently observed in the analysis of the 60 groups of homologous collinear blocks (Fig. 3). This suggests that genomic regions displaying the clade topology 2 correspond to the maternal contribution to the nuclear genome. We also measured the average nucleotide divergence of mitochondrial genes between the 3 apomictic *Meloidogyne* (*Mi*, *Mj* and *Ma*). On average, the inter-species nucleotide divergence was very low (0.17%) and ranged from 0 to 0.33%. In contrast, the average nucleotide divergence between the amphimicitic *M. hapla* and the 3 apomictics was 24.50 % and ranged from 24.42 to 24.58 %. Hence, mitochondrial phylogenetic analysis shows highly similar mitochondrial genomes between the three apomictic species.

### The duplicated genomes of asexual *Meloidogyne* allow potential functional divergence between gene copies

Apomictic *Meloidogyne* genomes are rich in gene copies with substantial divergence. To test whether this gene redundancy may result in a relaxation of selective pressure on the copies, potentially leading to functional divergence, we employed two different strategies. One raw approach simply based on pairwise computation of the ratios of rates of non-synonymous (Ka) vs. synonymous mutations (Ks); and a subtler approach using phylogenetic and statistical methods. To compute the average Ka / Ks ratios per pair of anchor homologous genes, we used the Nei-Gojobori method (Nei and Gojobori 1986) implemented in MCScanX. We found that 657 (8.8%) (*Mi*), 703 (20.5%) (*Mj*) and 2,082 (19.5%) (*Ma*) anchors gene pairs forming segmental duplications had a Ka / Ks ratio greater than 1, indicating possible positive selection (Fig. 5). In a second, phylogeny-based approach (Methods), we looked for signs of episodic diversifying selection (EDS) in duplicated genes within collinear regions shared by apomictic *Meloidogyne* genomes. We retrieved all homologous genes groups containing a gene copy for each of the three apomictic species and *M. hapla* and with at least one apomictic species having 2 or more copies. We found 1,735 such groups and used them to generate multi-gene alignments and their respective ML midpoint-rooted phylogenies. Using the random effects branch-sites model (Pond et al. 2011), we found 174 (*Mi*), 109 (*Mj*) and 207 (*Ma*) gene copies showing evidence of EDS (Supplementary Fig. S7 for an example) at the 0.05 confidence level (P_Holm: corrected for 9 tests using the Holm-Bonferroni procedure). Among these genes, 28 (*Mi*), 19 (*Mj*) and 55 (*Ma*) were also found to have Ka / Ks >1.

**Fig 5:**
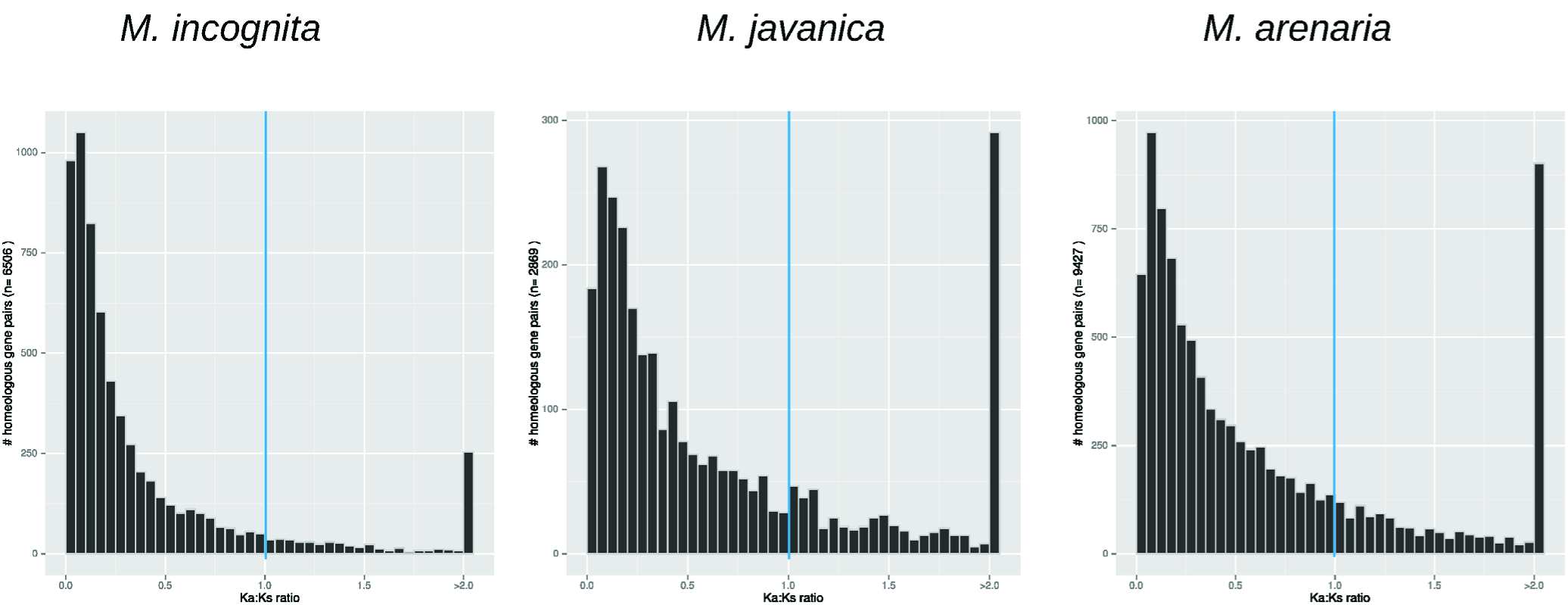
Distribution of the ratio of non-synonymous to synonymous substitutions rates per pair of anchor genes for *M. incognita, M. javanica* and *M. arenaria*. Histogram displaying the number of pairs of genes having a given Ka / Ks ratio in the three apomictic species *M. incognita*, *M. arenaria*, and *M. javanica*. Rates of non-synonymous (Ka) and synonymous (Ks) substitutions were calculated using the Nei-Gojobori method for each pair of anchor genes forming collinear blocks in the three apomictic species (see Methods). Pairs with a Ka / Ks ***>*** 1 indicate genes under positive selection.

To assess whether particular functional categories were affected by positive or episodic diversifying selection, we examined Pfam domains and gene ontology terms associated to these genes (Methods, Supplementary results S1). Overall, a large variety of Pfam domains and associated GO terms were identified among protein encoded by genes under positive selection (Ka / Ks >1) or subject to EDS, in the 3 apomictic species. As many as 399, 301 and 683 distinct Pfam domains, corresponding to 185, 169, and 318 distinct GO terms were found in proteins encoded by genes with Ka / Ks ratio >1 in *M. incognita*, *M. javanica* and *M. arenaria*, respectively (Supplementary Table S2). Similarly, we identified 124, 78 and 172 distinct Pfam domains, corresponding to 95, 56 and 112 distinct GO terms, in proteins encoded by genes under EDS in *M. incognita*, *M. javanica* and *M. arenaria*, respectively (Supplementary Table S3). Regardless of the dataset (Ka / Ks or EDS), few Pfam domains and GO terms were common to all the 3 apomictic species and the majority of them were related to enzymatic, binding and metabolic activities as well as transmembrane transport functions (Supplementary Tables S2, S3). At the Pfam domain level, there were more species-specific domains than domains shared by two or more species. Only 8 Pfam domains were identified as common to the 3 species in the EDS dataset and 5 of these domains were also common to the 3 species in the Ka / Ks dataset. More overlap was observed at the fine-grain GO level, with 86 GO terms shared by the 3 apomictic species in the Ka / Ks dataset, of which 18 were also shared by the 3 in the EDS dataset. At the coarse-grain level, with fewer and more generic GO-slim terms, much more overlap was found between the 3 apomictic species. The vast majority of GO-slim terms were shared by the 3 species both in the EDS and Ka / Ks datasets (Supplementary Tables S2, S3), and this observation was true for the 3 ontologies (Biological process, Molecular function and Cellular component). Overlap between the EDS and Ka / Ks datasets was also high as 26 of the 28 GO terms shared by the 3 apomictic in the EDS dataset were also shared by them in the Ka / Ks dataset (10/12 BP, 11/11 MF and 5/5 CC).

We identified significantly enriched Pfam domains as well as GO and GO-slim terms both in the Ka / Ks and EDS datasets as compared to the whole protein sets in each species (Supplementary Tables S2, S3). However, there was only little overlap at the Pfam domain level (only one commonly enriched term in the Ka / Ks dataset and one other in the EDS dataset). Furthermore, there was no overlap at all in enriched GO term either at the fine-grain or coarse-grain levels and this was true both for the Ka / Ks end EDS datasets. These observations suggest that most of the enriched protein domains and predicted functions associated to genes under positive or diversifying selection were different in each species rather than shared by the 3 asexuals.

### Asexual *Meloidogyne* genomes have a high number of genes and are rich in transposable elements

In the absence of a mechanism to control their proliferation, TE and repeats may invade the genomes of asexually reproducing animals (Arkhipova and Meselson 2005). We annotated transposable elements (TE) in the genomes of the 3 apomictic *Meloidogyne* as well as that of *M. hapla* and compared the TE content of the 4 genomes (Methods). Overall, we found that TE span 50.0, 50.8 and 50.8 % of the genome assemblies of the apomictic *M. incognita*, *M. javanica* and *M. arenaria*, respectively (Table 2). In comparison, TE occupy only 29.2 % of the genome assembly of the amphimictic species *M. hapla*. The genomes of the asexually reproducing *Meloidogyne* thus appear to be 1.7 times more rich in TE that that of the only sexual species whose genome is currently available. Consistent with this observation, Class I retroelements are on average 1.5 time more abundant in the apomictic species. Within Class I elements, DIRS-like (*Dictyostelium* intermediate repeat sequence) appear to have undergone a particular expansion in the asexuals as they are on average 5.5 times more abundant than in the sexual species. Class II DNA transposons are 1.9 times more abundant in the three apomictic species than in the *M. hapla* genome. Although Helitron occupy a comparable proportion in asexuals and in the sexual, all the other categories are more than 2 times more abundant in the asexuals. This includes Maverick-like and TIR (terminal inverted repeats) elements as well as “unclassified” TE that possessed characteristics of Class II elements but could not be further assigned to one family. The rest of the potential TE is in the “other” category, which gathers DNA fragments displaying contradictory features of both Class I and II elements. This category was also more abundant in the asexuals than in the sexual species (~1.8 times). This overall abundance of TE in asexual *Meloidogyne* has implications at the protein-coding level. While 27-30% of the protein-coding genes of asexual Meloidogyne are embedded within TE, only 17 % of *M. hapla* genes are within TE. Hence, TE abundance partly explains the higher number of genes observed in the asexual *Meloidogyne* (43,718-102,269 compared to 14,207 in *M. hapla*).

We tested whether the higher gene numbers observed in asexual *Meloidogyne* was homogeneously distributed along all protein domain families. We plotted the abundance of protein domains in *Mi*, *Mj* and *Ma* as a function of their abundance in *Mh* (Methods, Supplementary Fig. S8). The abundance of protein domains in *Mi*, *Mj* and *Ma* were all positively correlated to the abundance in *Mh* (R^2^= 0.92, R^2^= 0.89 and R^2^=0.87 for *Mi*, *Mj* and *Ma*, respectively). The slopes of the linear regressions were 3.06, 4.49 and 4.80 for *Mi*, *Mj* and *Ma*, respectively, suggesting that most of the protein domains are between 3 and 5 times more abundant in the three asexuals as compared to *M. hapla*. We compared the abundance of Pfam domains found in proteins encoded by genes present within TE and important for their own transposition activity (e.g. reverse transcriptase, integrase, transposase) in the 4 Meloidogyne species (Methods). We found that, on average, these domains were 3.4 to 9.8 times more abundant in asexual *Meloidogyne* than in *M. hapla* (Supplementary Table S4). For instance, rve (integrase core domain) is present in 205-689 copies in the three asexuals while it is found in only 59 copies in *M. hapla*. Similarly, the DDE_ 3 (DDE superfamily endonuclease) domain is absent in the *M. hapla* protein set while it is found in 13-61 copies in the three asexuals. This suggests that expansion of at least some families of TE might be in part responsible for the higher number of protein-coding genes in the asexuals.

## DISCUSSION

### Parthenogenetic Meloidogyne have large duplicated and rearranged genomes incompatible with meiosis

In a sexually reproducing diploid eukaryote, meiotic pairing and segregation require high sequence identity and collinearity between homologous chromosomes. Usually, sequencing the genome of a diploid sexual eukaryote involves performing repeated cycles of inbreeding to obtain lineages that are virtually homozygous at all loci. Genome assembly then results in collapsing all these virtually identical paternal and maternal variants into one single haploid reference sequence. This is exactly what is observed for the meiotic species *Meloidogyne hapla* which was assembled into a ~54Mb genome (Opperman et al. 2008). This genome assembly size is similar to reported experimental measures of haploid genome sizes (40-50 Mb) in the literature (Lapp and Triantaphyllou 1972; Pableo and Triantaphyllou 1989). Concordance of genome assembly size with experimental measures of haploid DNA content, associated to the absence of extensive duplications of genomic regions, indicate a canonical sexual diploid genome with highly homozygous parental and maternal haplotypes. The haploid chromosome number of *M. hapla* is n=16, similar to the putative ancestral haploid number of chromosomes (n=18) in *Meloidogyne* species (Triantaphyllou 1985; Castagnone-Sereno 2006; Castagnone-Sereno et al. 2013). Hence, we can hypothesize that the ancestral haploid genome size for a *Meloidogyne* is ~55-60 Mb with n= 16-18 chromosomes. The genome assembly sizes of the 3 mitotic *Meloidogyne* species we describe here range from ~180 Mb to ~260 Mb. These large genome sizes are confirmed by our flow cytometry measures of nuclear DNA content that suggest an even larger genome size of up to ~300 Mb for *M. javanica* and *M. arenaria* (Table 1). Hence, the genomes of apomictic *Meloidogyne* (~180-300 Mb) are 3 to 5 times higher than what can be considered as the ancestral haploid genome size for a *Meloidogyne* species (~55-60Mb, see above). Furthermore, Pfam domains are on average 3.06-4.80 times more abundant in apomictic *Meloidogyne* than in the haploid genome assembly of *M. hapla*. Finally, more than 90% of the protein-coding genes in the genomes of mitotic parthenogenetic *Meloidogyne* are present in at least two copies and a substantial proportion form collinear blocks of duplicated genome regions. Taken together, these results strongly suggest that the genomes of mitotic parthenogenetic *Meloidogyne* are polyploids (at least triploids). Similarity between genome assembly sizes and measures of total nuclear DNA content via flow cytometry suggest that most of the former paternal and maternal haplotypes have been separately assembled. This is further supported by the identification of duplicated collinear genome regions spanning several Mb and thousands of genes in the three apomictic genomes. These pairs of genomic regions have a similar average nucleotide divergence of ~8% within a species, confirming previous observations for the first genome assembly of *M. incognita* (Abad et al. 2008). Likewise, the per site synonymous substitution rate (Ks) of coding sequences of genes used as anchors to delineate duplicated genome regions had a very similar median of 0.1 for all three species. This homogeneity of average nucleotide divergence levels and of Ks between pairs of collinear regions for the 3 *Meloidogyne* species suggests that, for each species, the two copies have been separated for a comparable amount of time.

In *M. incognita* and *M. arenaria*, we observed collinear regions present in palindromic or tandem arrangement on a same scaffold. Such structures, similar to the ones observed in the ancient asexual bdelloid rotifer *A. vaga* (Flot et al., 2013), appear incompatible with conventional meiosis and confirm the strict asexual reproduction of these organisms. As to whether such structures are the cause or consequence of asexuality remains to be determined. Meiosis has never been observed in these asexually reproducing root-knot nematodes and, consequently, their nuclei are probably never present at a haploid stage. Palindromes or tandem blocks were not observed in the genome of the meiotic facultative parthenogenetic *M. hapla*, and because this genome presents high contiguity (the highest for a *Meloidogyne*), this most probably represents true absence. In *M. floridensis*, only 12 genes were found in one pair of duplicated regions (<0.1 % of genes), which is even less than in *M. hapla* (0.6 %). However, this is probably due to the highly fragmented and poorly assembled state (N50 = 3.5kb) of the *M. floridensis* genome sequence that prevented McScanX from identifying colinear duplicated regions. An improved genome assembly will be required to conclude on the presence of such blocks in the meiotic parthenogenetic too.

### Origin of the duplicated genome structure of apomictic Meloidogyne

The most frequently observed topologies in our phylogenomic analysis show that the duplicated genome regions tend to be more similar across different species than they are to their other copies within the same species. Furthermore, when duplicated regions form two or more clades in phylogenomic analysis, these clades more frequently present distinct topologies. Thus, collinear duplicated regions within a species have different origins and evolutionary histories and probably do not originate from common ancestral allelic regions that accumulated mutations separately (*i.e*. no Meselson-White effect). Contrasting with the high within-species divergence of duplicated blocks in the nuclear genome (avg. divergence ~8%), our mitochondrial phylogenetic analyzes shows that mitochondrial genes are almost identical in *M. incognita*, *M. arenaria* and *M. javanica* (avg. divergence ~0.17%). This confirms previous observation that these three species share virtually identical mtDNA markers (Hugall et al. 1999; Lunt 2008; Fargette et al. 2010) and suggests that *Mi*, *Mj*, and *Ma* share closely related or common maternal ancestors. The mitochondrial genome is expected to accumulate mutations faster than the nuclear genome (e.g. 100-1,000 times more rapidly than the nuclear genome in *C. elegans* (Denver et al. 2000; Denver et al. 2004; Denver et al. 2009)). Hence, the divergence time between the nuclear genome copies within a species is supposed to be much higher than the divergence time of the different species themselves, based on mitochondrial genomes. Inter-specific hybridization is the most likely hypothesis that could resolve at the same time the discrepancy between low mitochondrial and high nuclear divergence levels, the presence of diverged genomic regions in the genomes, and the observed topologies in the phylogenomic analysis (alternatives hypotheses are discussed in the Supplementary discussion S1).

In 1999, based on the presence of multiple divergent ITS nuclear ribosomal markers within apomictic *Meloidogyne* despite closely related mitochondrial markers between species, it was suggested that these species had undergone hybridization from a set of closely related females mating with more diverse paternal lineages (Hugall et al. 1999). More recently, a phylogenomic analysis of the first version of the *M. incognita* genome compared to the draft version of the *M. floridensis* genome also suggested that *M. incognita* was of hybrid origin (Lunt et al. 2014). However, the lack of contiguity of the *M. floridensis* genome prevented any analysis of conservation of collinear duplicated regions and thus all kinds of duplications were mixed together in this analysis (tandem, dispersed, segmental and WGD), blurring the overall signal. Furthermore, only one apomictic species was included in this analysis, preventing any comparison of within-species variation levels and between-species variation. Here, we were able to confirm at a whole genome scale that not only *M. incognita* but also *M. javanica* and *M. arenaria* most likely originated from multiple hybridization events with a same or closely related maternal donor lineage. Part of the collinear regions are present in more than 2 copies within a genome, suggesting that the three mitotic parthenogenetic *Meloidogyne* are at least hypo-triploid (*Mi*) and triploid or hypo-tetraploid (*Mj* and *Ma*) as a result of multiple independent hybridizations. Importantly, in most of the cases, the third copy of a gene was more similar to a cognate gene in another *Meloidogyne* species than to any of the other two copies found in the same species (Table 5). Thus, the third copies probably derive from a distinct hybridization event and not from species-specific duplication or inter-individual variation (unlikely in clonal lineages). This suggests a two-step hybridization process. First, homoploid hybridization (hybridization between two diploid progenitors without associated genome doubling) took place. Then, a second hybridization between an unreduced gamete of the homoploid hybrid with a reduced gamete of another sexual species led to the presence of three copies of nuclear genomes within a given species. It should be noted that unreduced gametes are frequently produced by inter-specific hybrids (Mason and Pires 2015). Following these successive hybridization events, genome rearrangements occurred and led to the synteny breakpoints and palindrome structures observed in the apomictic *Meloidogyne*. Whether this caused or followed the loss of meiosis remains to be established. It was recently proposed (Lunt et al. 2014) that *M. floridensis* is itself a hybrid and is one of the parents of *M. incognita*. According to this hypothesis, *M. incognita* would be a double hybrid of *M. floridensis* with another, yet undetermined parent. At the mitochondrial level, our phylogenetic analysis shows that the closest known relative of *M. incognita* is *M. floridensis*, a result also previously observed in previous phylogenetic analyses (Tigano et al. 2005; Adams et al. 2009; García and Sánchez-Puerta 2015). Furthermore, *M. floridensis* is the only known meiotic species in a clade otherwise only comprised of mitotic species (Castagnone-Sereno et al. 2013). It is thus tempting to speculate that *M. floridensis* could be the maternal donor of *M. incognita*, considering their high mitochondrial similarity. However, important additional supporting evidences are currently lacking. For instance, it is unknown whether the peculiar meiosis in *M. floridensis*, lacking a second division (Handoo et al. 2004), is compatible with amphimixis. Thus, another possibility would be that the donor might be another close relative of *M. floridensis* but not *M. floridensis* itself. Regardless of whether *M. floridensis* represents the maternal donor of *M. incognita*, the origins of *M. javanica* and *M. arenaria* appear to be distinct. Indeed, in mitochondrial and nuclear phylogenies, *M. floridensis* is not closely related to the three apomictic *Meloidogyne* but only to *M. incognita*.

### A high proportion of transposable elements in the genomes of apomictic Meloidogyne

Following loss of sexuality, it has been hypothesized that TE could invade the genomes in the absence of mechanism to control their proliferation (Dolgin and Charlesworth 2006). Alternatively, it has been suggested that the only asexual animals that survive are those that control TE multiplication in their genomes. Examples supporting these two contradictory hypotheses exist in the literature. In *Daphnia* arthropods, sexual reproduction seems to be correlated with a slower accumulation of TE in genomes (Schaack et al. 2010). Consistent with this, it has been shown that TE are more abundant in Wolbachia-induced asexual lineages of parasitoid wasps than in sexual lineages of the same species (Kraaijeveld et al. 2012). However, whether this is a consequence of sex loss or of Wolbachia infection remains to be clarified. In contrast, in the ancient asexual bdelloid rotifer *A. vaga*, TE occupy only 3% of the genome and while a high diversity of TE was found, they are generally present at very low copy numbers (Flot et al. 2013). This suggests that TE proliferation might be under control in this species. Recently, a comparison of the TE load in five sexual vs. asexual lineages of arthropods showed no evidence for TE accumulation in the asexuals (Bast et al. 2015). In the root-knot nematodes, we have found that TE occupy ~50% of the genomes of the three apomictic species while they occupy only 29% of the genome of the amphimictic species *M. hapla*. Although it appears that TE have proliferated in the genomes of the asexual *Meloidogyne*, whether this is a consequence of their mode of reproduction or a clade-specific feature will have to be clarified when further genome sequences of sexual and asexual root-knot nematodes from distinct clades will be available.

Regardless of their relation to the mode of reproduction, this abundance of TE in apomictic *Meloidogyne* might provide them with a mechanism of genomic plasticity in the absence of sexual recombination. Supporting this hypothesis, a transposon named Tm1 has been identified in apomictic *Meloidogyne* but no homolog with an intact transposase could be found in the sexually-reproducing relative *M. hapla* (Gross and Williamson 2011). Interestingly, the non-protein-coding gene Cg-1 whose deletion is associated to resistance-breaking strains of *M. javanica*, has been identified within one of these Tm1 transposons. Thus, TE possibly have a functional impact on these nematodes, including on their host plant range.

### Possible advantages conferred by hybridization and polyploidy

*M. incognita*, *M. javanica* and *M. arenaria* are exceptionally successful, globally distributed pathogens of diverse agricultural crops (Trudgill and Blok 2001; Perry et al. 2009). Intriguingly, their geographical distributions and hosts ranges are wider than those of their sexual relatives. Furthermore, in controlled condition, they are able to overcome plant resistance within a few generations (Castagnone-Sereno 2006). In the absence of sex and meiotic recombination that provide genomic plasticity allowing adaptation on the one hand and purge deleterious mutations on the other hand, their allopolyploid nature may endow them with several benefits that may explain their ecological success. First, polyploidy can provide the raw material for neo-and sub-functionalization of duplicated gene copies, resulting in novel genetic variation (Cuypers and Hogeweg 2014; Soltis et al. 2014). It has been shown in yeast that ploidy level is correlated to faster adaptation (Selmecki et al. 2015). Also, it has been suggested that polyploidy could mask deleterious recessive alleles (Madlung 2013) and limit their accumulation via gene conversion between homologous regions (Flot et al. 2013). Furthermore, allopolyploidy (due to hybridization) may provide transgressive phenotypes that surpass those of the parent species via novel genetic variation and heterosis (Madlung 2013; Mason and Pires 2015). Hybridization events, yielding plant pest species able to parasitize novel host plants that none of their progenitors could infest have already been described in insects (Schwarz et al. 2005) and in fungi (Menardo et al. 2016). It can be hypothesized that hybridizations could be involved in the larger host range of the asexual root-knot nematodes.

Here, we tested whether the presence of several divergent genomic copies in a same species, probably as a result of hybridization, could have functional consequences at the coding level. Hybridization brings together homeolog chromosomes and therefore orthologous gene copies within an individual. Among the many mechanisms reshuffling the genome, loss and degeneration of duplicated genes is expected to be important (Lynch and Conery 2000). Nonetheless, hybrid progeny may inherit orthologs with slightly different fitness due to adaptation of the parental species to different environments. Most likely, the hybrid inherits orthologs that had retained similar function and following functional redundancy, selective pressure on these genes may relax and drive them to different evolutionary trajectories (Ohno 1970; Lynch and Conery 2000). In some cases, the relaxation of selective pressure can allow emergence of new adaptive mutations. We have shown that ~8 to 20% of gene copies coming from the duplicated genomic regions harbor signs of positive selection. A diversity of Pfam domains and associated gene ontology terms were predicted in proteins encoded by positively selected genes. Although many terms and domains were related to enzymatic and other catalytic functions, there was a poor overlap between the 3 apomictic species, and different domains and functions were specifically enriched in positively selected genes in each species. These observations suggest that the functional consequences of the hybrid genome structure were different in each species. Thus, functional divergence between homeologous copies resulting from hybridization, appears to be rather unpredictable and specific to each species.

How an animal can survive without sexual reproduction and compete with its sexual relatives remain an evolutionary puzzle. Intriguingly, asexually-reproducing (apomictic) root-knot nematodes outcompete their sexual relatives as plant parasites of global economic impact. We have shown here that the genomes of these apomictic *Meloidogyne* are duplicated probably as a result of a complex series of hybridization events. The presence of large-scale duplicated and divergent genomic regions possibly promotes emergence of functional novelty between gene copies, following positive and diversifying selection. Furthermore, the TE-rich nature of their genomes might also foster genomic plasticity. Part of the intriguing success of apomictic root-knot nematodes could thus reside in their peculiar duplicated and TE-rich genomes. Whether this potential for genomic plasticity actually acts as a substitute for sexual reproduction, in generating the required functional malleability towards adaptation to a broad host spectrum and large geographic distribution, is something that future analyzes will have to further confirm.

## MATERIALS AND METHODS

### Genome assembly

DNA and RNA samples preparation and sequencing protocols are detailed in the Supplementary methods S2. Genome assemblies were performed in four steps, following the same procedure as for the genome of the bdelloid rotifer *A. vaga* (Flot et al. 2013): (i) assembly of 454 data into contigs, (ii) correction of the 454 contigs using Illumina data, (iii) scaffolding of 454 contigs and (iv) gap closing using Illumina data. For the first step, we used the multi-pass assembler MIRA (Chevreux et al. 1999) version 3.9.4 (normal mode, default options except the number of cycles) to generate contigs from the 454 genomic libraries (Supplementary Table S5). Moreover, Sanger reads of the *M. incognita* first draft genome sequence (Abad et al. 2008) were used to generate the current assembly. Twelve (*M. arenaria* and *M. javanica*) or sixteen (*M. incognita*) cycles were performed to separate a maximum of repeats and polymorphic regions. We subsequently used Illumina data to correct the homopolymer errors of the 454 contigs following a standard procedure (Aury et al. 2008). The corrected contigs were linked into scaffolds using the program SSPACE (Boetzer et al. 2011) with 454, Sanger and Illumina data. Finally, assemblies were gap-closed using GapCloser from the SOAPdenovo 2 package (Luo et al. 2012) with Illumina data. The statistics of the three genome assemblies are summarized in Supplementary Table S5. We assessed the completeness of the 3 genome assemblies by counting the number of Core Eukaryotic Gene (CEG) using CEGMA (Parra et al. 2007).

### Search for collapsed duplicated regions

To check whether some nearly identical duplicated genomic regions had been collapsed during the assembly (as previously observed in the *A. vaga* genome(Flot et al. 2013)), we aligned the Illumina PE-reads of each species against their respective genome assembly sequence. We computed the per base read coverage using BEDtools genomeCoverageBed (Quinlan and Hall 2010) and plotted the distribution of the per-base coverage depth. This clearly showed 2 peaks for the 3 species, one systematically at twice the coverage of the first peak (Supplementary Fig. S1). We calculated the number of bases with per base coverage comprised in the range of the second peak and summed it up to obtain the total size of the duplicated regions that had been collapsed during the assembly.

### Structural annotation

Predictions of protein-coding genes were performed using EuGene 4.1c (Foissac et al. 2008), optimized and tested for *M. incognita* on a dataset of 301 non-redundant full-length cDNAs. Translation starts and splice sites were predicted using SpliceMachine (Degroeve et al. 2005). Three datasets of *M. incognita* transcribed sequences were provided to EuGene to contribute to the prediction of gene models: i) Sanger ESTs (Genbank 20110419), ii) a dataset of two 454 transcriptomes obtained in our lab in a previous unpublished study, and iii) a dataset of nine Trinity (Haas et al. 2013) assemblies of RNAseq data, generated in this study. Transcribed sequences were aligned on the genome using GMAP (Wu and Watanabe 2005); spliced alignments spanning 80% of the transcript sequence length at a 90% identity cut-off were retained. Similarities to i) *C. elegans* release Wormpep221, ii) *G. pallida*, release 1.0 (Cotton et al. 2014), and iii) Swiss-Prot release December 2013 (excluding proteins similar to REPBASE (Jurka et al. 2005)) were searched using BLAST (Altschul et al. 1997) and provided to EuGene to contribute to gene modelling. The gene modelling algorithm used the standard parameters for the 4.1c version, except for the fact that i) the gene finding algorithm was applied on both strands independently allowing overlapping gene models, ii) non canonical GC/donor and AC/acceptor sites were allowed on the basis of transcriptional evidences, iii) a gene model was not allowed to span a gap (‘N’) longer than 1,000 nucleotides, iv) the minimum length of introns was set to 35 nucleotides, and v) the minimum CDS length cut-off was set to 150 nucleotides. For *M. arenaria* and *M. javanica*, the EuGene pipeline, with models and parameters tuned on *M. incognita*, was used to annotate both genomes. Two modifications were applied on the selection of reference datasets i) Swissprot (excluding proteins similar to REPBASE) and the proteome of *M. incognita* were used as reference proteomes ii) assemblies of *M. arenaria* and *M. javanica* RNAseq data were used as sources of transcription evidences.

We annotated ncRNAs using RNAmmer (Lagesen et al. 2007), tRNAscan-SE (Lowe and Eddy 1997), Rfam release 11 (Burge et al. 2013), and in house scripts to remove redundancy and consolidate results.

### Functional annotation

The predicted protein sequences of *M. incognita*, *M. javanica*, *M. arenaria* and *M. hapla* were scanned for the presence of Pfam protein domains using the program PfamScan (Finn et al. 2014) against the Pfam-A HMM domain library (release 27.0), using default thresholds and parameters. A gene ontology annotation was inferred from the Pfam protein domain annotation using the pfam2go mapping file maintained at the geneontology portal and generated from the InterPro2GO mapping (Mitchell et al. 2015). Gene ontology terms were also mapped on the generic GO-slim ontology using the GOSlimViewer utility developed as part of AgBase (McCarthy et al. 2006). The GO-slim annotations were split into three ontologies (biochemical function, cellular component and molecular function).

### Analysis of whole-genome duplication

The duplicated structures of *Meloidogyne* species wee estimated by detecting conserved blocks of duplicated genes. The protein sequences of each genome were initially self-blasted to determine a homologous relationship with an e-value threshold of 1e^−10^. Conserved blocks of duplicated genes were detected based on the gene locations in the genome using MCScanX (Wang et al. 2012) with default parameters. We required at least 3 collinear genes pairs (anchor genes) for MCScanX to form a block. Using the perl script, add_ka_and_ks_to_colinearity.pl included in the MCScanX package, we calculated Ks values for each homologous gene pairs between duplicated blocks. The median Ks value was considered to be a representative of the divergence between duplicated regions. We used custom python script to compute the pairwise nucleotide identity between collinear blocks for each species. Briefly, pairs of duplicated genomic regions were extracted according to the GFF3 positions of their first and last anchor genes. They were then aligned using NUCmer from MUMmer v3.23 (Kurtz et al. 2004) with default parameters. We then filtered out sub-alignment shorter than 50 nt (delta-filter −1 50) and summarized alignment using the dnadiff program from the MUMmer package. The average identity at the nucleotide level between duplicated regions was obtained from the output of dnadiff. Identity within coding and non-coding sequences was obtained by masking coding or non-coding sequences in each duplicated region before NUCmer alignment using BEDtools maskFastaFromBed v2.17.0 (Quinlan and Hall 2010).

To analyze synteny conservation between genomes, we concatenated all the inter-/intra-species BLAST hits (evalue threshold of 1e^−10^) of *M. incognita*, *M. javanica* and *M. arenaria* and *M. hapla* protein sequences and fed McScanX with this pooled BLAST result as well as with information on the location of the corresponding genes in the respective genomes, as recommended in the McScanX manual for multi-species comparisons. The *M. floridensis* genome had to be discarded from this comparative analysis because only one pair of regions composed of 12 genes block was detected for this genome preventing any large-scale analysis of conserved synteny in this species. We required at least 3 collinear genes pairs (anchor genes) for MCScanX to detect a block. We parsed the results of the collinearity analysis between genomes of *Meloidogyne* species (HTML files output by MCScanX) to extract anchor genes forming coinear regions conserved between *Meloidogye* species. We used those homologous anchor genes sets to perform phylogenomics analyses (see below).

### Determination of fragmentation bias

For each pair of duplicated regions, the genes present on each region were counted and the number of genes that were present in the ancestor of these two regions was calculated as the total number of genes on the two collinear regions minus the number of anchors genes pairs. We then compared the number of genes in each region to the number of estimated genes in the ancestral region to determine whether one region had lost significantly more genes than the other within a pair.

### Alignments, phylogenies and topologies searching

Multiple sequence alignments were performed using a custom python script. First, protein sequences were aligned using MUSCLE v3.8.31 (Edgar 2004a; Edgar 2004b). Second, protein alignments were back translated into codon alignment using PAL2NAL v.14 (Suyama et al. 2006) with the ‘nogap’ option. Third, codon alignments were trimmed using GBLOCKS (Castresana 2000) with default options. The fittest model of nucleotide evolution was searched using the function ModelTest as implemented in the R package phangorn (Schliep 2011). We then used phyml (Guindon et al. 2010) (-d nt -b -4 -m GTR -f e -t e -v e -a e -s BEST) to build maximum likelihood phylogenies with SH-like branches support on these pruned alignments. We rooted the phylogenies using the midpoint function of R package phangorn. For the rest of the analyses, we only retained the trees in which *M. hapla* held an outgroup position relative to the other *Meloidogyne* species in the midpoint-rooted topologies. Tree topologies were classified and counted using a custom R script. Phylogenetic tree figures were formatted and edited using EvolView (Zhang et al. 2012).

### Identifying gene copies subject to episodic diversifying selection

We performed tests of episodic diversifying selection (EDS) using the random effects branch-sites model (Pond et al. 2011) implemented in the HYPHY package (Pond et al. 2005). We looped the branchSiteREL.bf script over the 1,735 multi-sequence alignments and their respective ML midpoint rooted trees. Each alignment contained at least 1 anchor protein-coding gene for all three apomictic species and *M. hapla* and a duplicate in at least one asexual species. We chose the adaptive version of BSRE and allowed branch-site variation in synonymous rates. Branch with length less than 0.01 were not considered because ω rate classes cannot be inferred reliably for very small branches (<0.01).

### Experimental determination of nuclear DNA content

Flow cytometry was used to perform accurate measurement of cells DNA contents in the three apomictic *Meloidogyne (M. incognita, M. javanica* and *M. arenaria)* compared to internal standards with known genome sizes. *Caenorhabditis elegans* strain Bristol N2 (approximately 200 Mb at diploid state; (The C. elegans Sequencing Consortium 1998; Hillier et al. 2005) and *Drosophila melanogaster* strain Cantonese S. (approximately 350 Mb at diploid state; (Bosco et al. 2007; Gregory et al. 2013) were used as internal standards. Extraction of nuclei was performed as previously described (Perfus-Barbeoch et al. 2014). Briefly, for each *Meloidogyne* species about two hundred thousand stage 2 juveniles (J2s) were suspended in 2 mL lysis buffer (1mM KCl, 30 mM NaCl, 10 mM MgCl2, 0.2 mM EDTA, 30 mM Tris, 300 mM sucrose, 5 mM sodium butyrate, 0.1 mM PMSF, 0.5 mM DTT, 40 ⍰ Igepal), grinded for 10 min with a Dounce homogenizer and filtered through a 0.20 ⍰m nylon mesh. Subsequently, this 2 mL suspension was overlaid on top of 8 mL suspension buffer (same as lysis buffer except for sucrose, 1.2 M, and without Igepal) so that the tubes were ready for centrifugation (10000 rpm, 30 min, 4°C) to reduce the level of debris and to pellet nuclei. Supernatant was completely discarded and pelleted nuclei were re-suspended in suspension buffer. Then nuclei suspension was stained, at 37°C for 30 min, with 75 ⍰g/mL propidium iodide and 50 ⍰g/mL DNAse-free RNAse. The same nuclei extraction protocol was performed at the same time on the samples and on the two internal standards. Flow cytometry analyses were carried out using a LSRII / Fortessa (BD Biosciences) flow cytometer operated with FACSDiva v6.1.3 (BD Biosciences) software. Data were analyzed with Kaluza v1.2 software (Beckman Coulter) and cytograms exhibiting peaks for each phase of the cell cycle (G0/G1, S and G2/M) were obtained. Standards and samples were processed both alone and together. Only mean fluorescence intensity of the first peak (arbitrary units), corresponding to G0/G1 phase of the cell cycle of the cytograms, was taken into account to estimate DNA content. In this method (DeSalle et al. 2005; Hare and Johnston 2011), the amounts of DNA in the *Meloidogyne* samples were determined by interpolating the fluorescence signals generated from the standards using the following equation:

Meloidogyne DNA content (Mb) = (G0/G1_Meloidogyne sample_ x Standard DNA content)/ G0/G1_standard_. The estimated DNA contents of the *Meloidogyne* samples were calculated by averaging the values obtained from three biological replicates.

### Transposable Elements annotation and analysis

#### TE detection and Annotation

Repeat annotation was performed using the REPET pipelines TEdenovo and TEannot (Flutre et al. 2011). The TEdenovo pipeline was used to search for repeats in the contigs of the four Meloidogyne genomes pooled together (*M. incognita, M. javanica, M. arenaria, M. hapla)* to ensure that the annotation results were based on the same *de novo* predictions and were comparable between species. The high scoring segments pairs detection was performed by aligning all genomes against themselves using BLASTER (Quesneville et al. 2003) with the following parameters: identity >90%, High Scoring segments Pairs (HSP) length >100b & <20kb, e-value <1e^−300^. The LTRs retrotransposons were searched using LTRharvest (Ellinghaus et al. 2008) with the following parameters: LTR similarity >90% min LTR size >100bp & <1000bp. The repetitive HSPs identified from the BLASTER output were clustered using Recon (Bao and Eddy 2002), Grouper (Quesneville et al. 2003) and Piler (Edgar and Myers 2005). The predictions from LTRharvest were clustered using the MCL algorithm. The consensus for each cluster are obtained using MAP (Huang 1994) and classified with PASTEC Classifier (Hoede et al. 2014). The unclassified consensus sequences (noCAT) were filtered out to keep only consenus sequences built from at least 10 sequences in the cluster. The library of 10,535 consensus sequences obtained with TEdenovo was used for separate annotation of each genome using TEannot with the same parameters. The alignment of the reference consensus TE sequences on the genome was made using Blaster, CENSOR (Jurka et al. 1996) and RepeatMasker (http://www.repeatmasker.org/). The results of the three methods were concatenated and MATCHER (Quesneville et al. 2003) was used to remove overlapping HSPs and make connections with the “join” procedure. Using in-house perl script, we retrieved TE annotation with >80% sequence identity to consensus sequences & length >150 nucleotides.

#### TE-related Pfam domains

A published list of 124 Pfam domains associated with TE (Piriyapongsa et al. 2007) was combined with the list of TE-related Pfam domains included in the LTR Digest software (Steinbiss et al. 2009), resulting in a non-redundant list of 129 TE-related Pfam domains. The *Meloidogyne* species analyzed in the present article possessed 28 of these 129 TE-related domains. The compared abundance of these 28 domains in the 3 apomictic *Meloidogyne* is presented in Supplementary Table S4.

## ACKNOWLEDGEMENTS

This work was supported by the ANR program ANR-13-JSV7-0006 - ASEXEVOL, the INRA program AAP SPE 2011, as well as Université de Nice Sophia-Antipolis Postdoc program 2013-2014. We thank A. Robichon, JJ. Remy and DA. Fernandes De Abreu for providing *D. melanogaster* and *C. elegans* used as standards for flow cytometry experiments, C. Kim for help in counting tree topologies and formatting supplementary figures. We thank V. Jamilloux and the REPET team at INRA Versailles for help in installing and configuring REPET. The authors are grateful to the GenoToul bioinformatics platform Toulouse Midi-Pyrenees for providing computing resources. All the sequences generated in this research have been deposited in public database as detailed in the Supplementary note S1.

